# Risk optimization during ongoing movement: Insights from movement and gaze behavior in throwing

**DOI:** 10.1101/2024.12.10.627742

**Authors:** Stephan Zahno, Damian Beck, Ralf Kredel, André Klostermann, Ernst-Joachim Hossner

## Abstract

Handling motor noise is fundamental to successful sensorimotor behavior, especially in high-risk situations. Research using finger-pointing tasks shows that humans account for motor noise and costs of potential outcomes in movement planning. However, does this mechanism generalize to more complex movement tasks? Here, we investigate sensorimotor behavior under risk in throwing across three experiments with 20 participants each. Their task was to throw balls at a target circle, partially overlapped by a penalty circle. This task challenged participants to find strategies that trade off potential penalties and rewards. In the experiments, penalty magnitude and the distance between the circles were manipulated. We measured the location of their final gaze fixation before movement—as an indicator of their planned aiming point—and the ball’s impact location. Without penalty, the final gaze fixation and the ball’s impact location were both centered on the target. In the penalty condition, the location of the participants’ final gaze fixations and the ball’s impact shifted away from the penalty circle, with larger shifts for higher penalties and smaller distances. Interestingly, the shifts in the ball’s impact locations were not only larger (“more conservative”) but also closer to the statistically optimal (expected gain-maximizing) location compared to the fixated aim points. Movement trajectory analyses show that, in penalty conditions, the shifts away from the penalty zone increased until the final phases of the movement. These results suggest that risk evaluation is not completed in a pre-movement planning phase but is further optimized during movement execution.

**NEW & NOTEWORTHY:** We extend the study of sensorimotor behavior under risk from simple finger-pointing movements (Trommershäuser et al., 2008) to a complex throwing task in virtual reality. Our results suggest that, in complex sensorimotor behavior, risk evaluation of potential movements is not confined to a cognitive planning phase before movement but is optimized in action, with the motor system continuously biasing competing action options toward regions of higher expected rewards.

## INTRODUCTION

Dealing with noise in the sensory and motor system is a fundamental challenge in motor control [1-5]. Noise in motor commands limits the precision of our movements, unavoidably leading to errors. However, not all errors are equally costly. Trading off costs and rewards of potential movement outcomes is thus key to successful sensorimotor behavior, especially in high-risk situations. To illustrate with an extreme example: According to Swiss legend, the tyrant Gessler forced Wilhelm Tell to perform the horrible task of shooting an apple off the head of his son with a crossbow. The potential outcomes—either missing or hitting the apple or the boy—are clearly associated with different costs and rewards. Despite being the best crossbow shooter in town, Wilhelm Tell’s movement outcomes are still corrupted by variance and, therefore, risky in terms of hitting the boy. In statistical terms, selecting an aim point results in a probability distribution of potential outcomes. So, where should Wilhelm Tell aim?

Human sensorimotor behavior under risk has been studied in the lab, arguably with less vital consequences, using finger-pointing tasks. Trommershäuser and colleagues [6, 7] developed a task where participants received points by rapidly pointing to a green target circle on a screen while the target was partially overlapped by a red penalty circle. When accidentally hitting the penalty circle, participants were penalized by losing points. Due to severe time constraints, the pointing movements had to be executed with high speed, resulting in considerable outcome variance [8]. Thus, participants faced a similar dilemma as Wilhelm Tell and were challenged to trade off costs and rewards of potential movement outcomes.

In their seminal experiments, Trommershäuser et al. [6, 7] introduced a model of movement planning under risk based on statistical decision theory. According to the model, humans should estimate their motor variance to select aiming points that optimally trade off costs and rewards—i.e., maximize expected gains. The model of maximum expected gain is a normative model [9], meaning that it prescribes the optimal solution to the task, given biological constraints (here: participants’ motor variance). If the penalty circle overlaps the target circle on the left or right side, the model predicts that the optimal aim point horizontally shifts away from the center of the target circle as soon as the penalty is non-zero. In other words, participants should incorporate a “safety margin.” This horizontal shift should be larger with (1) higher penalty-to-reward ratios, (2) smaller distances between the centers of the target and penalty circle, and (3) higher motor variance. Trommershäuser et al. [6, 7] demonstrated that participants adjust their movements as predicted by the model, suggesting that they take into account their own motor variance and the costs of potential outcomes. This finding is intriguing, as it contrasts classical findings in cognitive decision-making tasks, where humans typically misinterpret probability information and fail to maximize expected gains [10], and suggests that humans are “more rational” in lower-level motor execution than they are in higher-level cognitive decision-making [11, 12]. Thus, studying sensorimotor behavior under risk offers a way to further our understanding of interactions between the cognitive and motor systems, which is currently regarded as a major challenge in the field of human behavior research [13].

How humans optimize risks in sensorimotor behavior has been extensively studied over the past two decades [6, 7, 11, 14-32]. The finding that humans take into account their own motor variance and (externally imposed) costs of potential outcomes to optimize their movements has become a key concept in motor control theory [33]. Conceptually, the idea is that humans acquire an internal model of their own motor variance and use probabilistic, predicted consequences of their movements to guide action. While studies have shown that human strategies are not always statistically optimal—e.g., when target-penalty configurations are asymmetric [19] or when the penalty level rapidly changes from trial to trial [23]— Trommershäuser et al.’s model has proven high explanatory power and provides a principled explanation of how humans deal with motor noise in sensorimotor behavior under risk [9, 34].

However, while handling risks in complex sensorimotor tasks is crucial in our daily lives, empirical evidence is limited to simple lab tasks [35, for a notable exception, see 36]. Arguably, the demands in these tasks, where participants remain seated and react to stimuli, typically using a chin rest to standardize the distance to the screen, considerably differ from the tasks we typically encounter in real-world situations. In recent years, going beyond simple lab tasks to test major theories under complex, naturalistic conditions in order to understand behaviors in situations they have evolved to function in, has been increasingly emphasized as a key challenge in psychology and neuroscience [13, 37-42]. Taking on this challenge is particularly relevant when aiming to transfer findings to applied settings, such as rehabilitation, transportation, surgery, or sports [35].

In this series of three experiments, we aim to bridge this gap by studying sensorimotor behavior under risk in a complex throwing task. Throwing balls or other objects, a complex multi-joint movement, is a fundamental sensorimotor skill that has evolved in humans since early hunter-gatherer societies and, in today’s society, remains essential in many sports [43]. We transferred the classical finger-pointing task by Trommershäuser and colleagues to a throwing task in virtual reality (VR). Participants throw balls at targets located on a wall at a distance of 3.2 meters. In this far-aiming task, considerable variance in movement outcomes arises naturally and does not need to be enforced by additional constraints in terms of a minimum execution speed. The experimental design was the same as in the Trommershäuser et al. studies. In Experiment 1 (Figure 1), participants received 100 points when hitting a green target circle, and they were instructed to collect as many points as possible across trials. The target was partially overlapped by a red penalty circle. We manipulated the amount of penalty and the distance between both circles, challenging participants to adapt their strategies and find solutions that trade off potential rewards and penalties.

**Figure 1.**
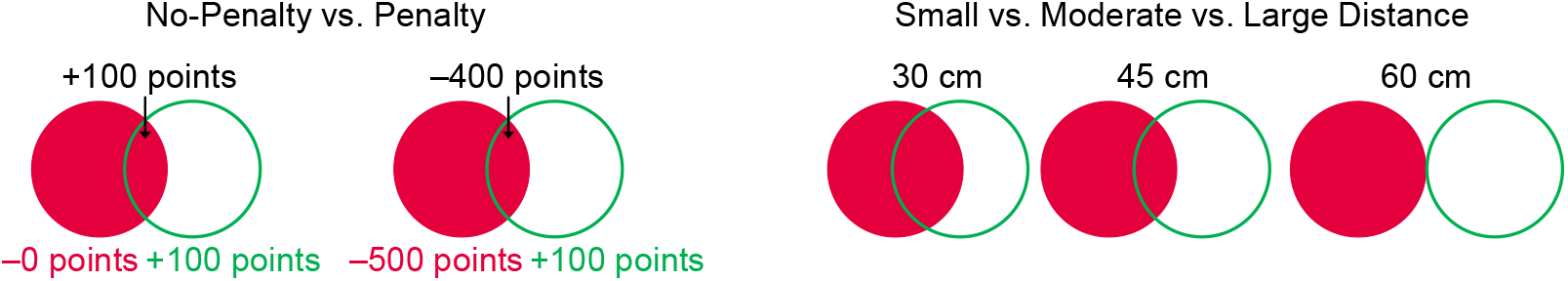
Conditions in Experiment 1. Participants’ task was to gain points by hitting the green target circle (+100). During the experiment, they faced 2 × 3 conditions: the penalty of accidentally hitting the red circle (0 vs. –500) and the distances (30 cm vs. 45 cm vs. 60 cm) between the centers of the target and penalty circle were manipulated. In Experiment 2, the colors of the penalty and target circle were swapped to exclude potential saliency effects. In Experiment 3, a second penalty condition (0 vs. –500 vs. –2000) was added to exclude fixation-location effects based on a purely geometrical optimization for information extraction.

Relevant differences between Trommershäuser et al.’s [11] and our task are the time constraints and the reactive versus active nature. In the Trommershäuser et al. experiments, participants had to place their index finger on a keyboard, and as soon as the target was displayed, they had 750 ms to react by lifting the finger and touching the screen. In contrast, the movement in our task is more self-paced. The target is displayed for around 5 s, allowing the participants to first localize the target, focus on it, and finally execute the throw without relevant time pressure. During the task, we measured participants’ gaze behavior and movement outcomes. As a first dependent variable, we assessed the location of participants’ final gaze fixation before movement execution, also known as the “quiet eye” in the far-aiming literature [44, 45]. Extensive research on gaze behavior shows that humans typically fixate their task goal prior to initiating goal-directed movements, suggesting that the location of their final fixation provides a valid indicator of participants’ planned aiming point before movement [46-52]. As a second dependent variable, we assessed the ball’s impact location relative to the target center as a measure of movement outcome. Notably, since the two dependent variables (final gaze fixation vs. ball’s impact location) relate to two different time points of interest (before and after the throw), the present throwing task under risk allows us to gain insights into risk-evaluation processes taking place before movement as compared to those evolving during movement execution.

In summary, the results of our three experiments show that participants adjust their behavior to changing spatial and penalty/reward conditions in a way that qualitatively aligns with the Trommershäuser et al. model. Interestingly, participants’ movement outcomes (the ball’s impact location) were consistently closer to the statistically optimal location than locations fixated before movement onset. This pattern suggests that evaluating risks of potential movements is not completed in the planning phase before movement but is further optimized during action execution. The motor system thus seems to play a key role in the deliberation process, biasing ongoing movements toward regions of higher expected reward. This interpretation is strengthened by an additional analysis of the movement trajectories, which shows that, in penalty conditions, the shifts away from the penalty zone (i.e., the “safety margin”) increase over the final phase of the throwing movement.

## MATERIALS AND METHODS

### Participants

In all three experiments, 20 young healthy subjects participated (Experiment 1: 10 females and 10 males, *M*_age_ = 21.42 years, *SD* = 1.40, 2 left-handers; Experiment 2: 7 females and 13 males, *M*_age_ = 20.90 years, *SD* = 2.28, 0 left-handers; Experiment 3: 4 females and 16 males, *M*_age_ = 21.55 years, *SD* = 1.69, 4 left-handers). All participants were naïve to the research questions and had normal or corrected-to-normal vision. The experiments were approved by the ethics committee of the Faculty of Human Science at the University of Bern (Approval Number: 2020-09-00003) and were conducted in accordance with the Declaration of Helsinki. All participants provided written informed consent.

### Apparatus

Participants performed underarm throws in a VR setup (Figure 2, left and video on GitHub: https://github.com/ispw-unibe-ch/risk_optimization_in_action). To display the virtual environment, we used the HTC VIVE Pro Eye head-mounted display (HMD) consisting of a Dual OLED 3 screen (2880 × 1600 px; 90 Hz refresh rate; 110° field of view), including an integrated mobile binocular eye tracker (120 Hz, 0.5°–1.1° within a FOV of 20°) (HTC Corporation, Taoyuan, China). We created the virtual environment using Unreal Engine 4 (Epic Games Inc., Potomac, MD, USA) and utilized SteamVR (Valve Cooperation, Bellevue, WA, USA), which provided the setup, room and eye-tracker calibration routines. Participants held a wireless HTC controller in their dominant hand, whose pose directly translated to a virtual 3-D representation of the participants’ hand. Participants could open and close the hand by pressing and releasing the trigger button, thus mimicking lifelike movements when grasping objects.

**Figure 2.**
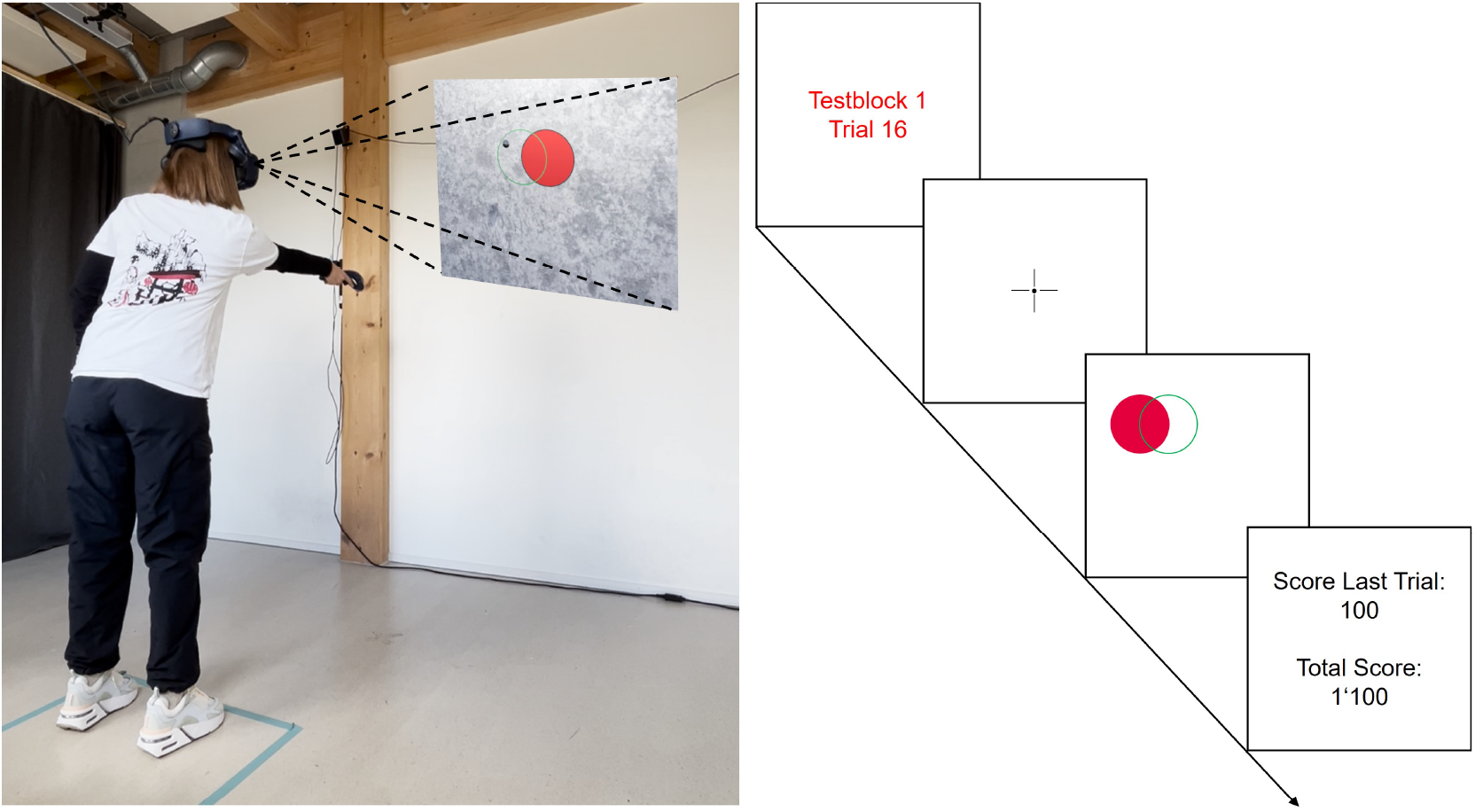
The virtual reality throwing task (left) and the trial procedure (right, showing an example of the last trial of a half-block).

In each trial, the virtual ball was initially positioned on the floor and automatically grasped by the virtual hand when holding the trigger button of the controller. When the trigger button was released, the virtual ball left the virtual hand. After ball release, ball trajectories were calculated within the game engine based on initial conditions (xyz-positions, xyz-velocities) and the classical Newtonian law of motion under gravity (no drag). The xyz-position when the ball hit the virtual wall was registered for each trial. In addition, the 3-D positions of the VIVE controller and the virtual ball, as well as binocular eye movements, were continuously collected with an average rate of 63.34 Hz (min: 45.45 Hz; max: 83.33 Hz). The fluctuations were due to the fact that the recording was realized within the game thread, which dynamically adjusts the refresh rate based on graphics-rendering requirements.

In the virtual environment, a line was displayed on the floor 3.2 meters in front of a gray wall on which the targets were displayed. The target and penalty circles both had a radius of 30 cm. Participants were instructed to stand behind this line when throwing the balls at the target. Before each half-block of 16 trials, the current penalty condition was displayed (either “penalty” in red or “no penalty” in green). Each trial lasted 10 s. As shown in Figure 2 (right), each trial started by displaying a text with the current trial number (when penalty, in red; when no-penalty, in green), followed by a fixation cross at the center of the wall, which disappeared 2.5 s after the start of the trial. After a short random delay (*M* = 0.78 s, min = 0.29, max = 1.24), the target and penalty circles were displayed on the wall. The circles disappeared 8 s after the start of the trial, meaning that participants had a time window of, on average, 4.72 s (min= 4.26 s, max = 5.21 s) for the throwing action, which in turn imposes no relevant time pressure for the execution of the movement. After the circles disappeared, the points achieved for the last trial were displayed for 1.5 s. After each half-block of 16 trials, the current total accumulated score was additionally displayed.

### Experimental design

Our experiments followed the same design as developed by Trommershäuser et al. [6, 7]. Participants came to our lab on two days, with exactly a one-week break in between. Day 1 was a training day. Participants started with 16 warm-up throws before having 12 blocks of 32 trials, resulting in a total of 384 throws. On day 1, all trials were in the no-penalty condition—that is, the penalty circle was visually displayed but irrelevant. Participants were instructed to collect as many points as possible by hitting the target circle (100 points per hit). The functions of day 1 were to get used to the VR environment, learn the throwing task, and—without mentioning this in the instruction—acquire knowledge about their own motor variance.

Data to test our hypotheses were collected on day 2. To this end, participants were given the same task goal—namely, to collect as many points as possible by hitting the target (100 points per hit). However, as shown in Figure 1, we now manipulated the consequences of hitting the penalty circle (no penalty = 0 points vs. penalty = –500 points), as well as the distance between the centers of both circles (30 cm vs. 45 cm vs. 60 cm). When participants hit the overlapping zone between the target and the penalty circle, reward and penalty were combined (no penalty = 100 points vs. penalty = –400 points). Participants were also informed that day 2 was a competition, with the best three participants being rewarded (vouchers for a bookstore: first prize = 50 CHF, second prize = 25 CHF, third prize = 10 CHF). As on day 1, participants started with 16 warm-up throws before having 12 blocks of 32 trials. In the total of 384 throws, all conditions (2 penalties x 3 distances) were provided in a blocked fashion in two blocks of 32 trials, which in turn were presented in a quasi-random order.

Potential confounding effects of the block ordering and the position of the penalty and target circles were addressed in the protocol. The conditions changed from block to block. Half of the participants were assigned to a protocol with blocks changing in a sequence from 30 cm to 45 cm to 60 cm and alternating between penalty and no-penalty blocks. For the other half of the participants, the protocol was exactly mirrored to balance potential sequence effects. Within each block of 32 trials, the penalty circle was placed to the right of the target circle in half of the trials and to the left in the other half. To avoid preplanned movements to a specific location at the wall before the target appeared, the spatial location of the target varied from trial to trial along four horizontal (15 cm and 45 cm to the left/right from the fixation cross) and four vertical positions (15 cm and 45 cm above/below the fixation cross) in a quasi-random order. That is, the target locations were distributed in a quite large area of 90 cm × 90 cm on the wall, forcing participants to adjust their movements at every trial. Within each block of 32 trials, each of the resulting 16 target positions appeared twice: once with the penalty to the right and once to the left of the target circle.

To ensure that the participants understood the instructions and the attribution of points, three test questions were asked after the instructions. To avoid fatigue effects, a five-minute break was included after every four blocks. Each experimental session lasted about 1.5 hours.

### Measures

#### Final gaze fixation

We assessed the location of participants’ final gaze fixation before movement initiation as an indicator of their planned aim point before execution. Specifically, for the underarm throwing movement, we used the location of the last fixation before the hand’s forward swing [cf. 53]. As a dependent measure, we calculated the directional horizontal shift of the fixation location away from the target center (in cm) and defined directionality so that higher values represent larger shifts away from the penalty zone (i.e., independent of the relative left/right position of the penalty circle in respect to the target circle).

To obtain the location of the final gaze fixation, we conducted the following steps. First, we detected fixations on the virtual wall using the dispersion-based algorithm by Nyström and Holmqvist [54], which classifies a fixation as soon as the point of gaze becomes stable within a circular area of 1.2° of visual angle for at least 120 ms. Second, using the controller data, we detected the timeframe of the reversal point in the throwing movement (i.e., the last frame before the start of the forward swing). Third, we excluded fixations before target onset (i.e., before the target was displayed). Finally, we selected the last fixation with a fixation onset before the initiation of the hand’s forward swing. For each participant, the mean horizontal shift in the final gaze fixation was aggregated for each of the six penalty x-distance conditions. Trials were excluded if no fixation on the virtual wall was detected in the relevant timeframe (Experiment 1: 1.1% of trials, Experiment 2: 0.5% of trials, Experiment 3: 0.6% of trials). There was no systematic effect of the condition on the proportion of invalid trials.

#### Ball’s impact location

We assessed the ball’s impact location as a measure of the movement outcome. As for the final gaze-fixation location, the dependent measure was the mean directional horizontal shifts of the ball’s impact locations away from the target center (in cm), higher values again representing larger shifts away from the penalty zone. Throwing trials that did not hit the wall (i.e., hitting the floor or the ceiling) were classified as invalid and therefore excluded (Experiment 1: 15.7% of trials, Experiment 2: 15.0% of trials, Experiment 3: 9.5% of trials), again without any systematic effect of the condition on the proportion of invalid trials.

### Modeling of the statistically optimal aim point

To compare participants’ strategies (i.e., their horizontal shifts away from the penalty region) to an optimal shift, we applied the model by Trommershäuser et al. [6, 7]. This model estimates the optimal shift that maximizes expected gains given the endpoint variance of participants’ throws (i.e., their motor variance). Our Python code is available on GitHub (https://github.com/ispw-unibe-ch/risk_optimization_in_action).

Each throw can result in one of four possible outcomes: the ball lands in the reward region (*R*_*reward*_) associated with a positive gain (*G*_*reward*_); in the penalty region (*R*_*penalty*_) associated with a negative gain (*G*_*penalty*_); in both regions simultaneously, resulting in a combined gain; or outside both the target and penalty regions, which yields zero points. Due to inherent noise in motor execution, participants exhibit variance in the ball’s landing positions (*σ*). While participants can choose an intended aim point, the movement outcome is probabilistic.

Each horizontal shift (x) is associated with an expected gain (Γ), formalized as:

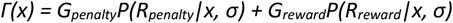

where *P(R*_*i*_|*x, σ)* represents the probability of hitting a region *i* given a horizontal shift *x* and the subject’s motor outcome variance *σ*. To calculate the expected gain, the probabilities are multiplied by the associated gain (*G*_*i*_) for each region. The optimal shift is the one that maximizes the subject’s expected gain.

Following Trommershäuser et al. [6, 7] as well as more recent studies [e.g., 22, 55], we used Monte Carlo simulations to find the optimal horizontal shift for each participant in each condition (30 vs. 45 vs. 60). The only free parameter in this model is the participants’ endpoint variance. First, we computed an endpoint covariance matrix for each participant based on their experimental data across conditions. We used these covariance matrices to model each participant’s endpoint variance in both the x and y dimensions with a bivariate Gaussian distribution. Next, we defined a one-dimensional grid of potential aim points, ranging from x = 0 cm to x = 60 cm (target radius = 30 cm), in 1 cm increments, and simulated 1,000,000 endpoints around each possible aim point to estimate, for each aim point, the probabilities of different outcomes (*R*_*penalty*,_ *R*_*reward*,_ or outside). By multiplying these probabilities with the gains associated with each region, we generated a landscape of expected gains for each horizontal shift. The optimal horizontal shift for each individual participant in each condition is the one with the highest expected gain across the landscape.

### Inferential statistical analysis

To test whether participants change their strategies—that is, their mean horizontal shifts in the final gaze fixation or ball’s impact location—as a function of conditions, we conducted 2 (penalty: 0 vs. –500) x 3 (distance: 30 vs. 45 vs. 60) ANOVAs with repeated measures. Additionally, to examine systematic differences between participants’ fixated aim points before movement initiation and their movement outcomes, we also conducted 2 × 3 ANOVAs with repeated measures on the differences between final gaze fixation and the ball’s impact location. We report η_p_^2^ as effect sizes and a priori set the α level to 0.05. The sample size of 20 was a priori defined to allow sufficient power for detecting medium-sized interaction effects in the 2 × 3 conditions design (α = .05, 1-β = .80, *f* = 0.25). The assumptions of normality and sphericity were evaluated prior to conducting the repeated-measures ANOVA. Normality was assessed using the Shapiro–Wilk test, while sphericity was examined with Mauchly’s test. Appropriate corrections (Greenhouse–Geisser) were applied when the assumption of sphericity was violated. All tests were performed using the Python package pingouin [56]. Furthermore, we qualitatively compared participants’ actual horizontal shifts with the shifts predicted by the maximum expected gain model. With the exception that, due to an additional penalty condition (penalty: 0 vs. –500 vs. –2000), 3 × 3 ANOVAs were required in Experiment 3, the same analyses were conducted for Experiments 1, 2, and 3.

## RESULTS

### Experiment 1

#### Shifts in the final gaze-fixation location and the ball’s impact location qualitatively align with the maximum expected gain model

To examine whether participants adapt their fixated aim points prior to movement as a function of penalty and distance between target and penalty circle, we compared the locations of their mean final gaze fixations (horizontal shift away from the target center) across the penalty (0 vs. –500) and distance conditions (30 vs. 45 vs. 60) (Figure 3, left). As predicted, mean final gaze fixations are centered on the target in the no-penalty condition (green), whereas in the penalty condition (red), the mean final gaze fixations shift increasingly away from the target center with smaller distances between the circles (i.e., more overlap), indicated by a significant penalty x-distance interaction (*F*[2, 38] = 43.20, *p* < .001, *η*_*p*_^*2*^ = .69). The shifts in fixated aim points suggest that, already before movement, participants consider a “safety margin” and adjust their aim point in a way that is in qualitative agreement with the Trommershäuser et al. [6] model.

**Figure 3.**
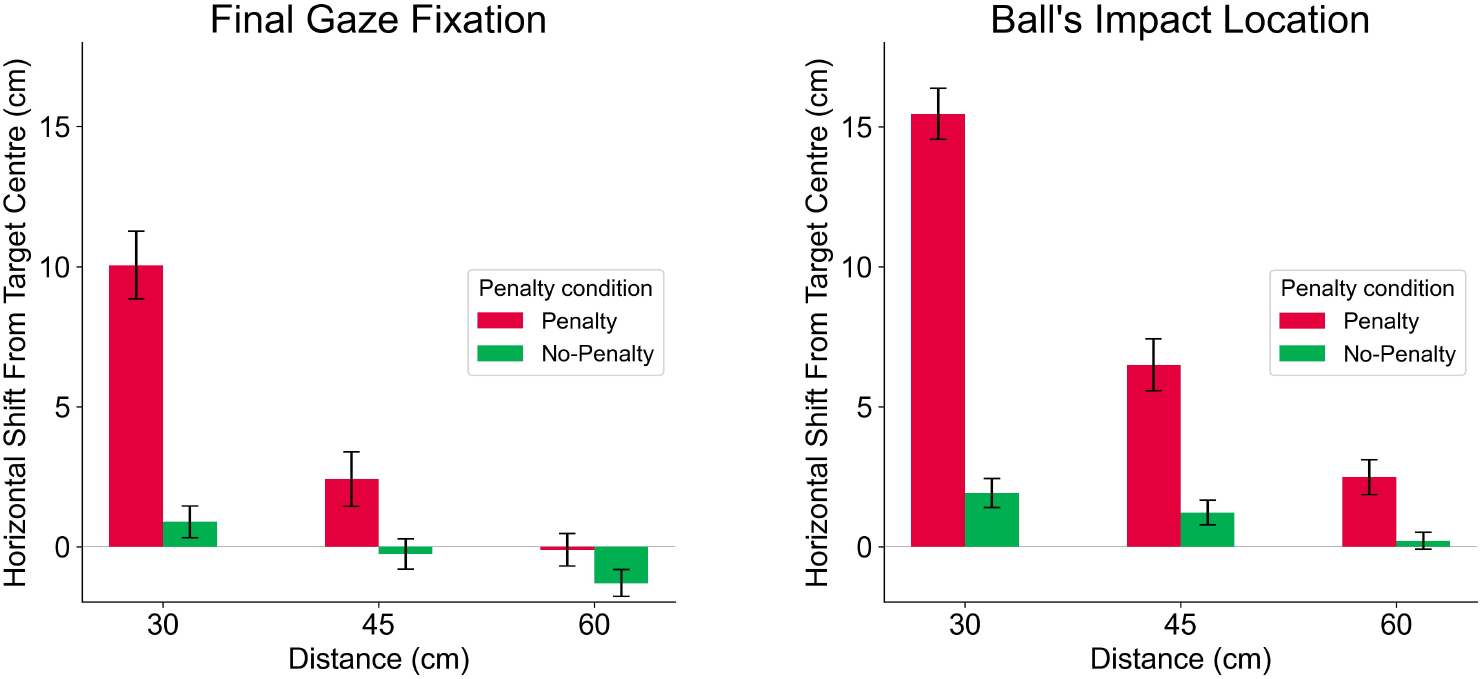
Horizontal shift in participants’ final gaze fixation (left) and in ball’s impact location (right) as a function of penalty (0 vs. –500) and distance between the centers of the circles (30 cm vs. 45 cm vs. 60 cm) in Experiment 1. Error bars represent standard error (SE).

For the ball’s impact location, we observe the same pattern (Figure 3, right), underpinned by a significant penalty x distance interaction (*F*[2, 38] = 106.49, *p* < .001, *η*_*p*_^*2*^ = .85). Consistent with the findings by Trommershäuser et al. [6, 7], these shifts were found already in the first test block of day 2 and did not further increase during the experiment. This suggests that participants acquired an estimate of their own motor variance during the training day and were able to immediately make use of it in the decision-making task under risk. A comparison of individual behaviors with quantitative predictions of the maximum expected gain model reveals an overall satisfying fit (Figure 4). However, the comparison also shows that a considerable number of participants tend to shift insufficiently, as indicated by data points on the left side of the dashed bisection line. The observation that some participants seem to underestimate their motor variance (or deliberately take too much risk) aligns well with previous findings using the classic pointing task [7] as well as timing-related visuomotor tasks [31, 57].

**Figure 4.**
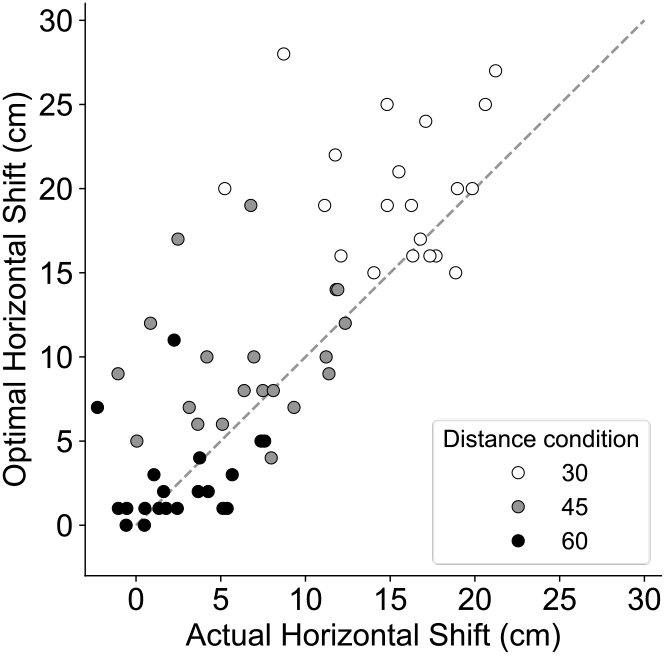
Participants’ actual horizontal shifts of mean ball’s impact location (x-axis) compared to the optimal horizontal shifts to maximize expected gains given their motor variance (y-axis) as a function of distance between the centers of the circles (30 cm vs. 45 cm vs. 60 cm) in Experiment 1.

Comparing the results of the final gaze-fixation location and the ball’s impact location leads to a notable observation: The horizontal shift (i.e., the “safety margin”) is more pronounced in the ball’s impact location than in the final gaze fixation (Figure 3, left vs. right). To follow up, we analyzed these differences as a function of the experimental conditions. In the no-penalty condition, differences between the fixation and impact locations were small and independent of the distance condition, whereas in the penalty condition, the differences increase with smaller distances, underpinned by a significant interaction effect (*F*[2, 38] = 4.44, *p* = .019, *η*_*p*_^*2*^ = .19). This suggests that, as soon as penalties are involved, subjects’ movements (i.e., the ball’s impact location) are more “conservative” than the fixated aim point before movement (i.e., the final gaze-fixation location).

When—instead of differences—analyzing the extent to which the ball’s impact location is related to the location of the final gaze fixation, we found a small correlation between both variables at a single-trial level (*r* = .20, *p* < .001). However, when averaging the final gaze fixations and ball’s impact locations for each participant and condition, the correlation is large (*r* = .77, *p* < .001). This suggests that while the final gaze fixation is indicative of where participants aim, the movement outcomes are not fully determined by the fixated aim point. In turn, the remaining variance can be attributed to motor noise and possibly, at least partly, also to corrections in later phases of movement execution.

#### Movement outcomes are closer to optimal than fixated aim points before movement

The experimental design also allows to compare the horizontal shift variables in terms of their relative position in the expected gain landscape (Figure 5). As expected for the no-penalty conditions, the mean fixated aim points (blue), ball’s impact locations (yellow), and optimal aim points (orange) horizontally align independently of the distance condition (Figure 5, top). In contrast, small shifts in the final gaze fixations are apparent in the penalty conditions, suggesting that participants do already consider a “safety margin” before movement onset. The shift in the movement outcome (yellow) is closer to the optimum (orange), which implies that the trade-off between penalty and rewards is further optimized during the unfolding movement (Figure 5, middle and bottom). Paired t-tests (two-sided) indicate that the horizontal difference between the ball’s impact location and the optimal aim point is significantly smaller than the difference between the final gaze-fixation location and the optimal aim point for both the distance-30 condition (*t*[19] = -4.95, *p* < .001, *d* = 0.88) and the distance-45 condition (*t*[19] = -4.23, *p* < .001, *d* = 0.73). In addition, the expected gain plots show that the participants do, on average, not fully shift their movements as much as they should, given their motor variance and potential penalties. However, they do reach a region in the landscape where a further shift would only marginally increase their expected gain. Together, these data suggest that (a) the horizontal shift in the ball’s impact location is not only more “conservative” than the fixated aim point but also closer to the statistically optimal (expected gain-maximizing) shift; (b) the trade-off between penalty and rewards seems to be further optimized during the unfolding movement; and (c) in doing so, participants reach a region in the expected-gain landscape that is close to optimal.

**Figure 5.**
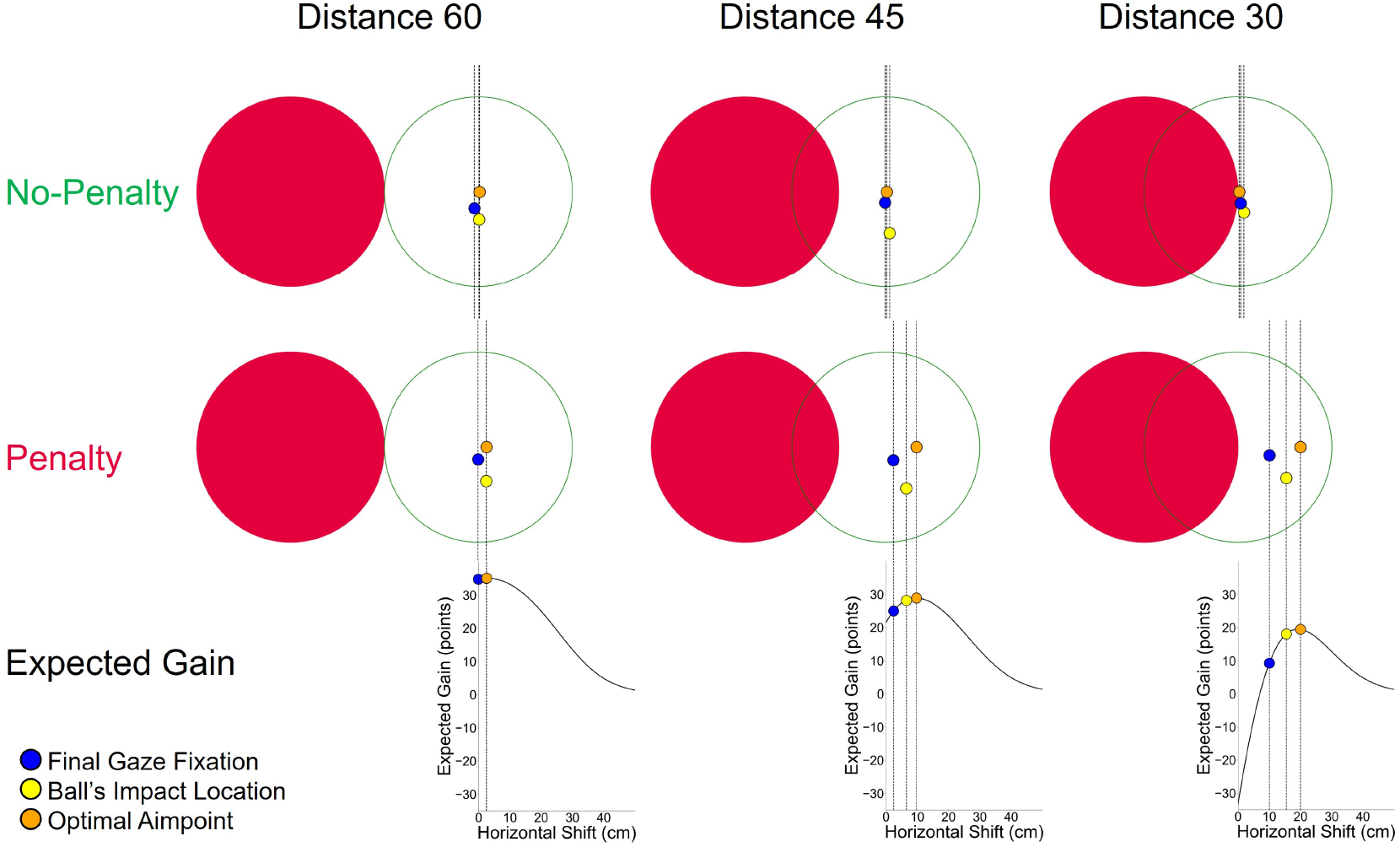
The final gaze fixations, ball’s impact locations, and optimal aim points as a function of distance between the centers of the circles in the no-penalty (top) and penalty conditions (middle), in comparison to respective expected gains for the penalty condition (bottom) in Experiment 1.

### Experiment 2

A potential alternative explanation for the results in Experiment 1 could be that the visual saliency of the red penalty circle attracts participants’ gaze bottom-up, resulting in a systematic difference between the final gaze fixation and the ball’s impact location. To examine whether our data could alternatively be explained by saliency effects, we conducted Experiment 2 with 20 new participants. The only difference between Experiment 1 and 2 is that the color of the target and penalty circles were swapped. We thus instructed the new participants that they gain 100 points when hitting the filled, red target circle and are penalized when hitting the green circle.

#### Differences between the final gaze fixation and the ball’s impact location cannot be explained by a saliency effect

Ruling out that our finding can be ascribed to a mere saliency effect, Experiment 2 fully replicates the pattern obtained in Experiment 1, with remarkably similar effect sizes (Figure 6). The interaction effects of the 2 (penalty: 0 vs. –500) x 3 (distance: 30 vs. 45 vs. 60) ANOVA with repeated measures are significant on both the final gaze-fixation location (*F*[2, 38] = 35.77, *p* < .001, *η*_*p*_^*2*^ = .65) and the ball’s impact location (*F*[2, 38] = 82.46, *p* < .001, *η*_*p*_^*2*^ = .81), as well as on the difference between the final gaze fixation and the ball’s impact location (*F*[2, 38] = 5.25, *p* = .01, *η*_*p*_^*2*^ = .22).

**Figure 6.**
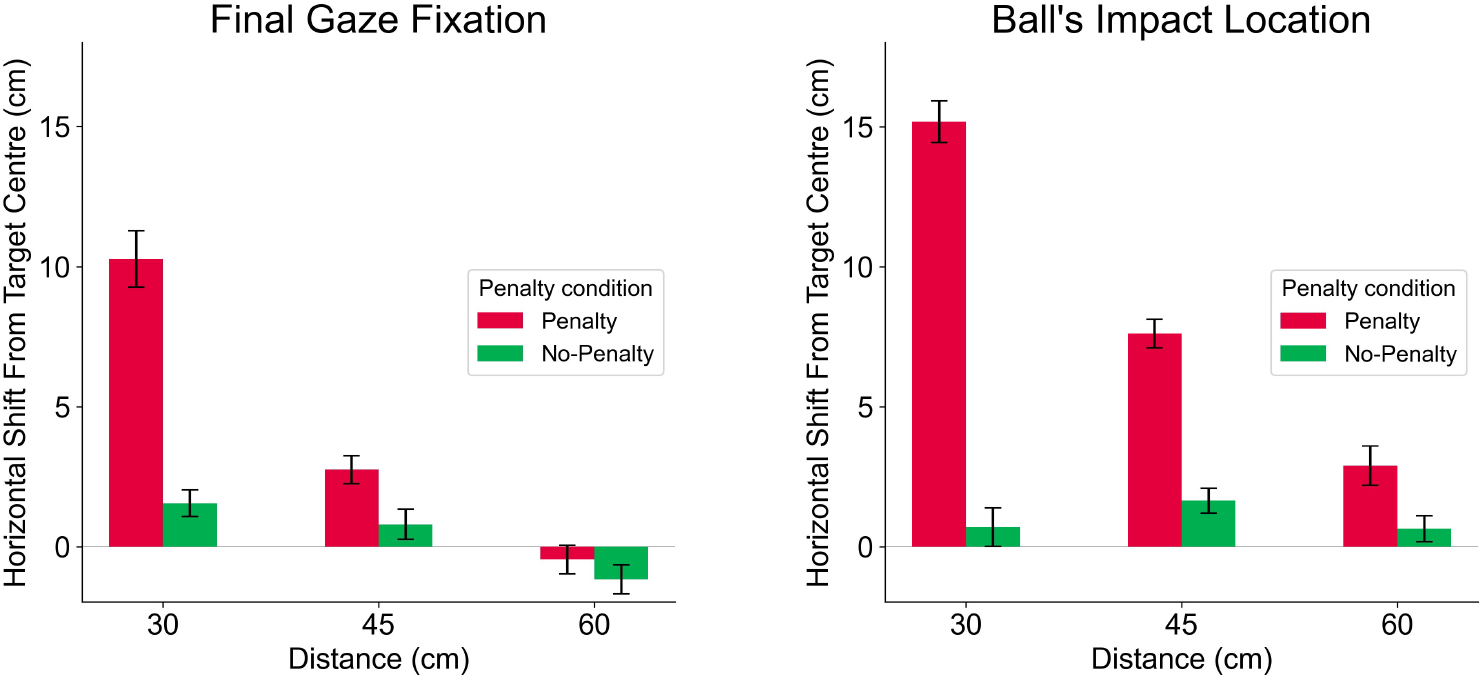
Horizontal shift in participants’ final gaze fixation (left) and ball’s impact location (right) as a function of penalty (0 vs. –500) and distances between the centers of the circles (30 cm vs. 45 cm vs. 60 cm) in Experiment 2. Error bars represent standard error (SE).

### Experiment 3

A second potential alternative explanation is that the final gaze fixation reflects a purely geometrically convenient location for information extraction rather than a planned aim point. When penalties are involved, a gaze anchoring between the edges of the penalty and the target, approximately as found in Experiments 1 and 2, appears optimal to fulfill this function. To test this alternative explanation, we conducted Experiment 3 with 20 new participants. The only difference between Experiments 1 and 3 is that we now added a third penalty condition, namely penalty –2,000 in addition to penalty –500 and penalty 0 (resulting in a 3 × 3 conditions design with 32 trials for each condition). If the final gaze fixation is purely optimized for information extraction and is not related to participants’ intended aim, no differences in the final fixation location between penalty –500 vs. –2,000 should be expected.

#### Final gaze-fixation location cannot be explained by a purely geometrical optimization for information extraction

Experiment 3 not only replicated Experiments 1 and 2, indicated by significant interaction effects for the final gaze-fixation location (*F*[4, 76] = 21.14, *p* < .001, *η*_*p*_^*2*^ = .53) and the ball’s impact location (*F*[4, 76] = 28.48, *p* < .001, *η*_*p*_^*2*^ = .60), but it also revealed that participants do shift their gaze more pronouncedly with increasing penalties (Figure 7). Using a priori contrasts and one-sided t-tests, the difference between penalty –500 and –2,000 in the final gaze fixation is significant for the distance-30 condition (*t*[19] = 1.99, *p* = .03 *d* = 0.44). This suggests that the final gaze fixation not just reflects a geometrically optimal location for information extraction but does provide an indicator for participants’ planned aim point before movement execution.

**Figure 7.**
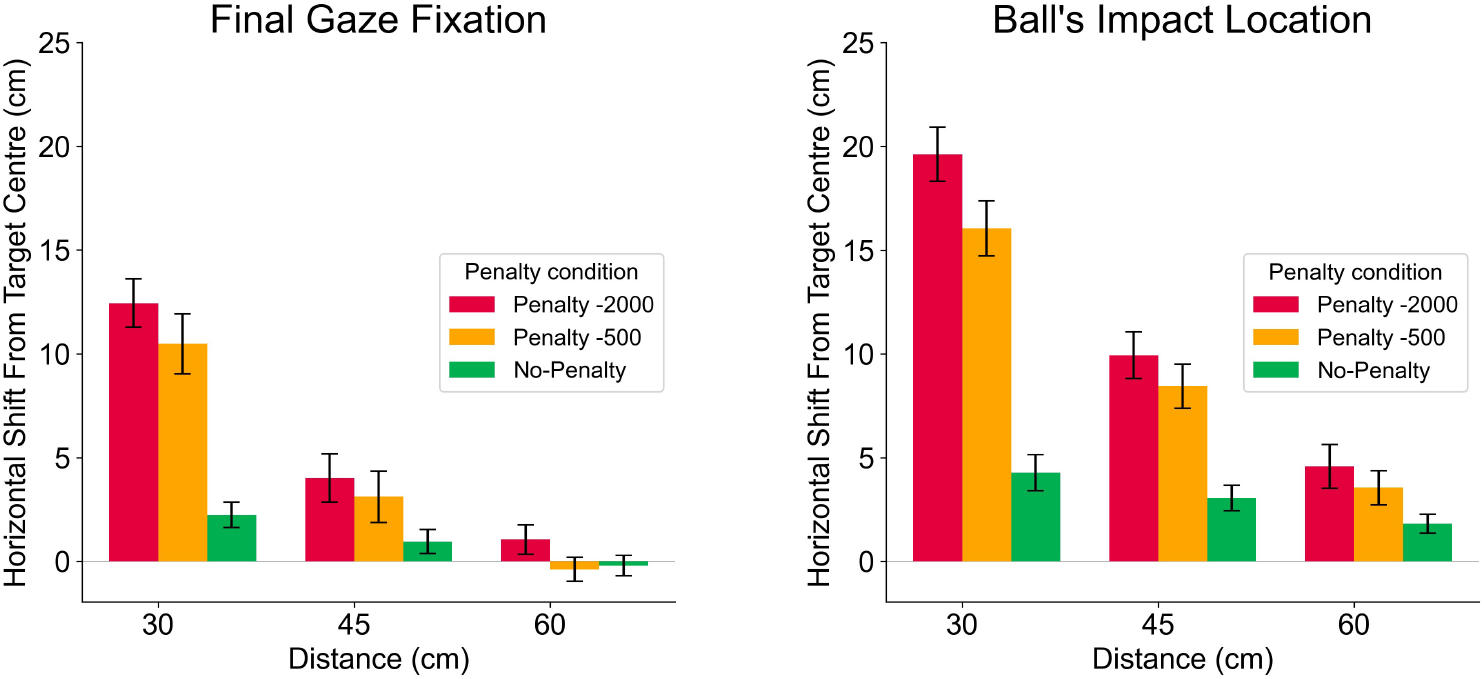
Horizontal shift in participants’ final gaze fixation (left) and ball’s impact location (right) as a function of penalty (0 vs. –500 vs. –2,000) and distance between the centers of the circles (30 cm vs. 45 cm vs. 60 cm) in Experiment 3. Error bars represent standard error (SE).

### Trajectory analysis

After having ruled out alternative explanations of our result pattern, we additionally examined whether we could find further support for our interpretation that the trading off between penalty and rewards is not completed in a planning phase but is further optimized during the unfolding movement. To this end, we fell back on the kinematic data from the hand controller and analyzed the movement direction in the horizontal plane during the final 200 ms of the forward swing before ball release. Since the total duration of the forward swing (i.e., the time interval from reversal point to release) was, on average, 379 ms (*SD* = 87 ms), varying from 200 ms to 756 ms, the time window of 200 ms before release represents a phase of forward movement in all trials of the experiment. For this time window, available movement trajectory data was cleaned by first removing extreme outliers (subsequent data points leading to unrealistic directional changes of more than 45°) and replacing them with cubically interpolated values. Subsequently, data were filtered using Savitzky-Golay (window length = 5) and resampled at 45 Hz to obtain constant time steps between each data point. Based on the cleaned data, we aggregated mean trajectories across participants for each condition. Trajectories where the penalty circle was on the right side were mirrored. Based on the movement direction derived from consecutive frames and linear extrapolation, we finally projected where the ball would have landed when released 1–8 timeframes before the actual release. Figure 8 depicts a single exemplary trial (top-down perspective) to illustrate the approach.

**Figure 8.**
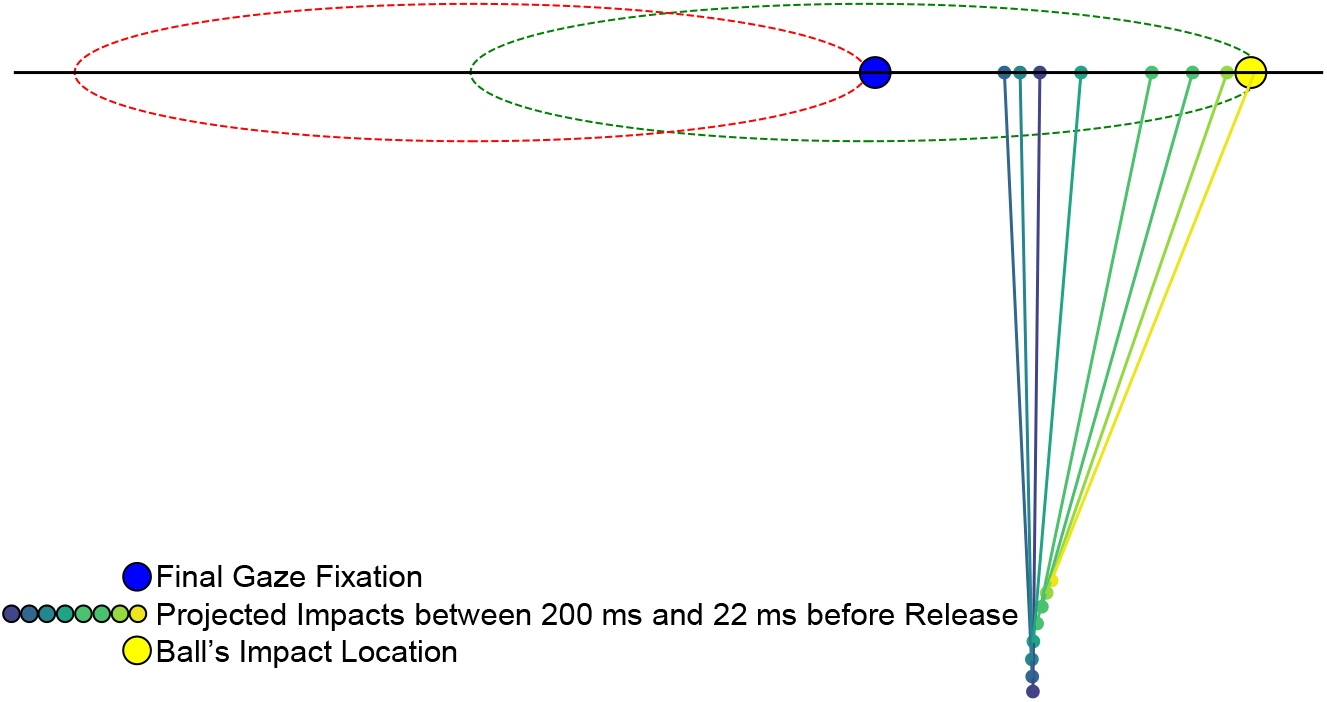
Exemplary trajectory data of one single trial (2-D, top-down perspective). The projected impact points over time were calculated based on the movement direction derived from two consecutive data points of the throwing movement and linear extrapolation.

Figure 9 illustrates, for the whole sample of Experiment 1, how the projected impact locations shift over the last 200 ms of the forward swing before ball release (dark blue to brighter yellow dots representing the temporal unfolding). This analysis based on aggregated trajectories shows that, on average, the movement direction is optimized until ball release. Specifically, in the penalty conditions, the shifts away from the penalty zone and toward the optimal aim point increase until the ball is released. Such a shift over time is not found in the no-penalty condition, suggesting it is related to optimizing risks in ongoing action.

**Figure 9.**
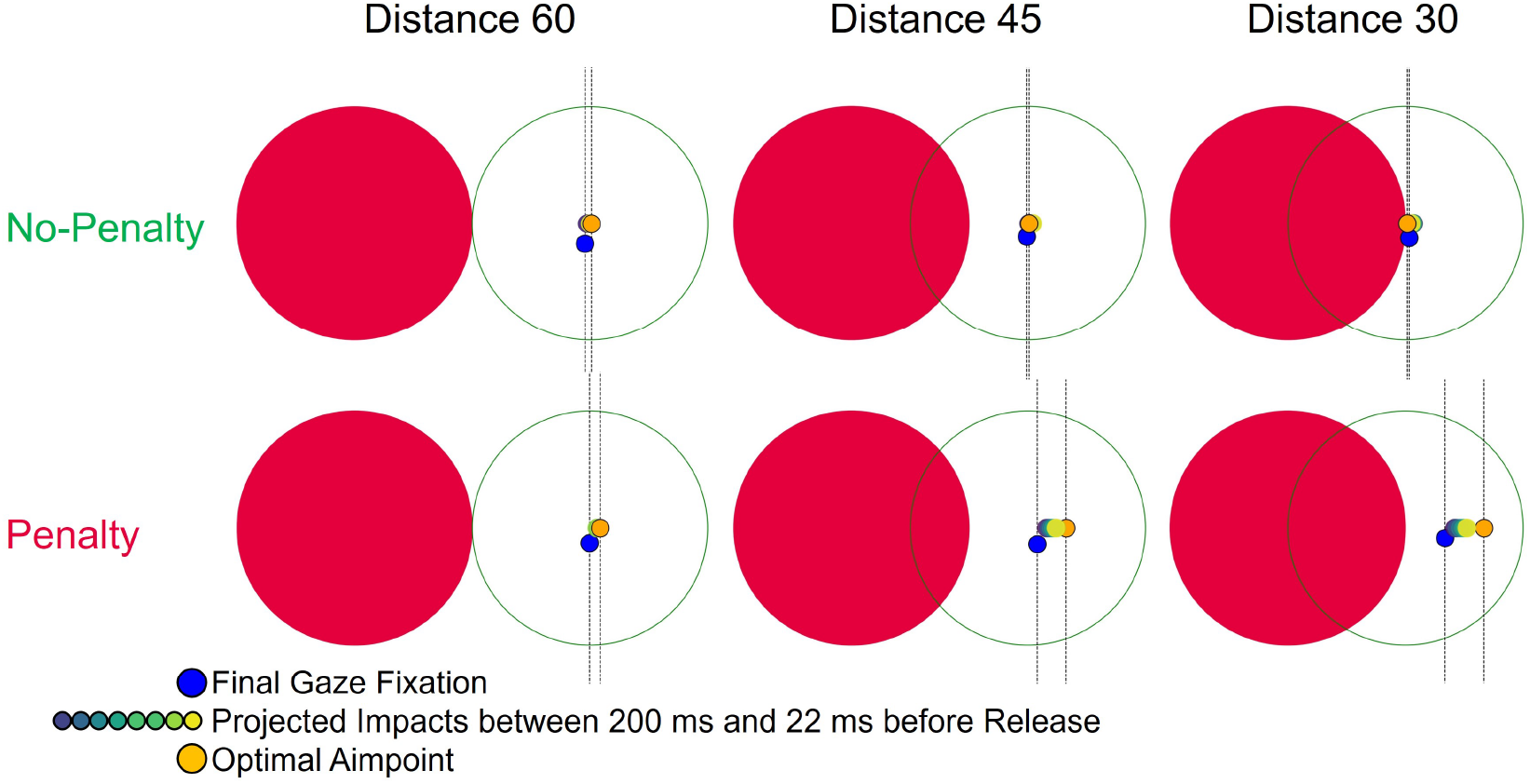
Final gaze fixations, projected impact locations over the last 200 ms of the forward swing before ball release, and optimal aim points, as a function of the distance between the centers of the circles in the no-penalty (top) and penalty conditions (bottom) in Experiment 1.

It should be added that, on top of the increasing shifts over time in the penalty conditions, our participants had a general tendency to perform throws curved to the left. The degree of curvature varies with the target’s position, indicating that biomechanical factors constraining the throwing movement also matter, and not all curves indicate online corrections. However, in penalty conditions, the left curvature is more pronounced when the penalty circle is on the right side and less pronounced when the penalty circle is on the left side, both compared to the no-penalty condition. Taken together, the data overall align with our hypothesis that humans optimize risk evaluations in action.

## DISCUSSION: An Online Risk Optimization Hypothesis

The aim of this study was to gain insights into human strategies in complex sensorimotor behavior under risk. To this end, we adapted the classical pointing task by Trommershäuser et al. [11] to a sports-like throwing task, allowing us to capture participants’ strategies at different time points: their aiming locations fixated before movement initiation, the ball’s impact location on the wall, and in between the projected throwing directions over the unfolding movement. We found that participants use strategies that are qualitatively consistent with predictions from the maximum expected gain model [11]. In no-penalty conditions, both participants’ final gaze fixations and the ball’s impact locations were, on average, aligned and centered on the target. With penalties, participants incorporated a “safety margin” by horizontally shifting their fixated aim points and movement outcomes away from the penalty circle. Extending the findings of Trommershäuser et al. [11], we observed that the shifts in penalty conditions were not only larger in the ball’s impact locations than in the fixated aim points (i.e., “more conservative”) but also closer to the statistically optimal locations. These findings suggest that risk evaluation of potential movement outcomes is not finalized in a pre-movement planning phase but is instead further optimized during movement execution, a concept we would like to refer to as *Online Risk Optimization*. Supporting this interpretation, an additional trajectory analysis shows that, in penalty conditions, the horizontal shifts away from the penalty zone increased over the final phase of the throwing movement.

Our study strengthens and extends the body of research on sensorimotor decision-making under risk [6, 7, 11, 14-25, 31, 32] by showing that statistical decision theory provides a principled framework for complex tasks. This aligns with recent calls to advance human sensorimotor-control research from simple lab tasks to more naturalistic tasks [37, 38, 40]. Understanding how humans deal with risks in sensorimotor behavior in real-world situations is pivotal for many applied fields, such as rehabilitation, transportation, surgery, and sports. For example, a slalom ski racer must continuously trade off the risk of getting too close to a pole and straddling a gate against taking a safer but slower trajectory. For sports, our work thus aims to provide a theoretical framework for understanding how athletes master such challenges and, ultimately, to support coaches to improve practice.

Beyond extending models from simple lab tasks to more complex sensorimotor demands, our research offers novel insights that may not have been revealed in classic paradigms. While Trommershäuser and colleagues [11] focused—by the nature of their pointing task—on movement planning, our data suggests that risk evaluation is not limited to a (cognitive) planning phase prior to movement onset but continues to be optimized during (motor) execution. This insight fits well with current models of sensorimotor behavior, such as the *Affordance Competition Hypothesis* [58], which posits that decision-making and movement control are much more intertwined than classically conceptualized [e.g., 59]. The idea that both processes operate in a parallel rather than in a serial manner with deliberation processes extending into movement execution is backed up by mounting behavioral [13, 60-65] and neurophysiological [66-69] evidence. According to the *Affordance Competition Hypothesis*, multiple potential actions are specified in parallel, continuously compete against each other, and are biased by the desirability of their predicted outcomes [70].

In this view, it is functionally plausible that participants anchor their gaze to an aiming location before movement, while specifying a set of potential options for action in parallel. When no penalties are involved, this location is centered on the target, and there is no reason why it should systematically shift during action. With penalties, participants already take into account costs and an internal estimate of their own motor variance when setting this initial aim point; however, they do so in a suboptimal way, possibly due to an underestimation of their motor variance. During subsequent active motor execution, participants appear to have an improved estimate of their own motor variance, presumably based on a continuously updated forward modeling of action consequences [71]. These refined predictions of the motor system would then continuously bias the competition between action options toward regions of higher expected gain until the point that matters most: ball release.

In conclusion, we propose an *Online Risk Optimization Hypothesis* which states that risk evaluation is not completed in a planning phase before movement but is instead further optimized during motor execution. Certainly, further and more direct testing of this hypothesis in future studies is required. Besides the systematic manipulation of noise, which can be implemented by altering the physics in our VR task, we see three approaches to test the online aspect of our hypothesis: designing experiments that include (a) the modification of relevant information after movement onset (e.g., penalty-reward ratio), (b) the manipulation of the richness of sensory feedback during movement execution, and (c) the comparison of conditions allowing for online corrections (such as in the current experiment) versus conditions where adjustments after movement onset are impossible (an aim-point selection task). Such experiments are planned in our lab, and we would be delighted if this work motivates other labs to join us in this inquiry.

## DATA AVAILABILITY

Data used for analysis and figure generation with the corresponding code (Jupyter Notebooks) as well as a video of the experimental task are available on GitHub (https://github.com/ispw-unibe-ch/risk_optimization_in_action)

## ACKNOWLEDGMENTS

We would like to thank Nina Hämmerli for her support in data collection, Martin Widmer and Fabio Bücheler for technical support, and Heather Neyedli for sharing an earlier version of the Monte Carlo Simulation code.

## DISCLOSURES

No conflict of interest.

## AUTHOR CONTRIBUTIONS

S.Z., A.K., E.-J.H. designed the experiments; R.K. developed the experimental setup; S.Z. and D.B. led the experiments; S.Z. analyzed and interpreted the data in collaboration with D.B., A.K., R.K., E.-J.H.; S.Z. drafted the manuscript; S.Z. and E.-J.H. edited and revised the manuscript.

## Notes

### Competing Interest Statement

The authors have declared no competing interest.

